# Data aggregation and mechanistic modeling enable dose-response analysis of SARS-CoV-1 in non-human primates

**DOI:** 10.64898/2026.06.01.729195

**Authors:** Philip C. Lee, Celine E. Snedden, Dylan H. Morris, James O. Lloyd-Smith

## Abstract

Dose-response modeling provides estimates of infectious and lethal doses, which can be used to inform control and prevention measures. Unfortunately, data from experimental challenge studies, which are needed to perform dose-response modeling, are often sparse. For example, non-human primate (NHP) challenge studies tend to have small samples sizes and little dose variation, often with only one or two dose levels per study. Thus, it is infeasible to apply traditional dose-response modeling approaches to data from single NHP studies. To address this challenge, we developed a mechanistic Bayesian model that aggregates and analyzes NHP pathogen load data across multiple studies. Our model links dose-infectivity to pathogen kinetics, which allows us to estimate the infectious dose and evaluate dose effects on within-host viral kinetics simultaneously. With this model, we obtained the first-ever ID_50_ estimate for SARS-CoV-1 in NHPs using data compiled from six NHP challenge studies. Our work demonstrates the value in reusing previous data from animal experiments. Our modeling framework can be applied to other pathogens, enabling robust dose-response inference when individual challenge studies are inconclusive.

**Author summary:** Dose-response models are used to estimate pathogen doses needed to cause infection in humans, so they are useful for informing outbreak control policies. Unfortunately, performing dose-response modeling can be difficult due to limitations in the available data. If the pathogen causes significant risk of severe disease or death in humans, then controlled human infections cannot be performed. Additionally, experimental challenge studies of relevant animal models, such as non-human primates (NHPs), often have small sample sizes and limited dose ranges, which make dose-response modeling unfeasible using data from single studies. We developed an approach to aggregate data across multiple challenge studies to enable dose-response modeling in the absence of dose-response experiments. We applied our approach to data from six NHP challenge studies to perform the first-ever dose-response analysis of SARS-CoV-1 in NHPs. Our approach also included a mechanistic, mathematical model of within-host pathogen kinetics, which allowed us to assess the effect of SARS-CoV-1 dosage on patterns of viral RNA shedding. The framework we developed can be readily applied to other host-pathogen systems, and the mechanistic components of our model contribute to a growing movement towards understanding dose effects beyond simple infectivity.

## Introduction

Dose-response models are a key component of the quantitative microbial risk assessment (QMRA) framework. They are used to quantify the risk of a given response (e.g., infection or death) to given doses of pathogen [1]. These models are often used to generate estimates of infectious and lethal doses, which are the doses required to cause infection or mortality in an individual with a given probability (e.g., the ID_50_ is the dose that produces infection in 50% of hosts). Obtaining estimates of the infectious dose in humans, such as the ID_50_, is especially useful for informing risk analyses and control policies. For example, a low ID_50_ may indicate the potential for high human-to-human transmissibility, and it could suggest the need for control measures that reduce even brief exposures.

Unfortunately, obtaining reliable estimates for infectious and lethal doses in humans is typically challenging due to limitations of the available data and the types of experiments that can be performed. In cases where the pathogen of interest causes significant risk of severe disease or death in humans (which makes controlled human infections unethical), dose-response model estimates must rely primarily on data from animal experiments, often using rodents [2]. Although these small animal model experiments are more feasible, and it is possible to obtain larger sample sizes which bolster statistical analyses, there are clearly inherent limitations when trying to extrapolate results from these data to human infections [3]. For example, early in the COVID-19 pandemic the best estimate of the infectious dose for SARS-CoV-2 in humans came from a study of SARS-CoV-1 in transgenic mice (ID_50_ of 280 PFU) [4], and the knowledge gap on human infectious dose was cited repeatedly as a limitation in guiding control policy [5, 6].

A natural route to obtain more human-relevant infectious dose estimates would be to apply dose-response modeling approaches to data from animal models that are more closely related to humans, such as non-human primates (NHPs). However, working with data from NHP experiments is also challenging. Due to important ethical and logistical considerations, NHP experiments tend to have small sample sizes, which makes it difficult to perform quantitative analyses using data obtained from a single experiment. Furthermore, many NHP studies are designed to investigate disease pathogenesis or evaluate countermeasures rather than estimate dose-response relationships, and thus they typically administer only one or two dose levels. Although individual studies often span a narrow dose range, inoculation doses can vary substantially across studies. This creates an opportunity to enable dose-response modeling by aggregating data from multiple studies to construct a synthetic dose series, while simultaneously increasing the total number of observations available for quantitative analyses.

Combining data across NHP challenge studies introduces several complications. Methodological differences across studies must be accounted for during data aggregation. The richness of data collected from these experiments further complicates analysis. Following inoculation with a pathogen of interest, animals are commonly sampled at multiple time points, across multiple anatomical sites (e.g., nasal/oral swabs, rectal swabs, blood samples), and with multiple assays (e.g., immunological measurements, PCR results, culture results). This can make it difficult to define which individuals are “infected” and “non-infected,” a classification that is required for traditional binary dose-response models. At the same time, these data present valuable opportunities to explore dose effects beyond simple infectivity. Individual-level variation in infection responses is well-documented, and a continuum of outcomes is often more realistic than a binary classification. For example, there is evidence that the within-host pathogen loads can determine host infectiousness [7, 8]. Additionally, the relationship between exposure dose and within-host pathogen kinetics has been a topic of previous study [6, 9]. For NHP experiments that report quantitative measurements for within-host samples (e.g., cycle threshold (C_T_) or copy number for quantitative PCR results, or other pathogen load values), there is the opportunity to investigate potential dose effects on within-host pathogen kinetics.

Here, we developed an approach to aggregate and reuse data from experimental challenge studies of NHPs to perform dose-response modeling. We used Bayesian computational methods to flexibly integrate data across multiple studies and to account for possible between-study biases. To do so, we created a mechanistic within-host kinetics model that links dose-infectivity to viral load trajectories, enabling simultaneous estimation of infectious dose and evaluation of potential dose effects on pathogen kinetics. This framework addresses the limitations posed by small sample sizes and narrow dose ranges in individual studies while making fuller use of quantitative virological data. We applied our approach to previously published virological data compiled from six SARS-CoV-1 NHP challenge studies. Our meta-analytic dataset comprised 39 individuals inoculated with doses ranging from 10^3^ to 10^6.9^ TCID_50_. We fit our model to observed upper respiratory tract (URT) viral load measurements in our dataset to produce the first-ever ID_50_ estimate for SARS-CoV-1 in NHPs. The within-host kinetics component of our model also allowed us to evaluate the effect of inoculation dose on the magnitude (initial viral load) and timing (day post infection of peak viral load) of the viral load trajectories in successful infections. This work provides a dose-response analysis of SARS-CoV-1 in a more human-relevant animal model, develops a framework for investigating pathogen dose effects beyond infectivity, and demonstrates the value of reusing data from previous animal challenge experiments.

## Results

### Data and Model Overview

We conducted a comprehensive literature search for SARS-CoV-1 NHP challenge studies. We found that the most common inoculation procedure was intranasal inoculation using nasal drops. For consistency, we only included studies in our analysis that performed intranasal inoculation of NHPs and reported viral load data from URT swabs (since intranasal inoculation with drops primarily leads to URT infections) [2]. We identified six studies that reported this type of data (**Table 1**). From these studies, we extracted 180 URT viral load observations from 39 individual NHPs. Two individuals from Lawler et al. (2006) were inoculated with SARS-CoV-1 via a combined intranasal and conjunctival route (50/50 split of the inoculum between the nose and conjunctiva). These individuals were included in our analysis because fluid desposited in the eye largely drains to the nasal cavity via the nasolacrimal duct [10], and thus these individuals should experience responses similar to individuals inoculated solely through the nose. Overall, the individuals in our dataset represented four exposure doses (10^3^, 10^5^, 10^6^, 10^6.9^ TCID_50_), two NHP species (rhesus and cynomolgus macaques), both sexes, and two age groups (juveniles and adults; **Fig 1**).

**Table 1.** Summary of studies included in the dose-response and kinetics analysis. “No. of Pos.” counts the number of individuals with at least one non-negative RT-PCR measurement at any time point. “No. of Obs.” is the total number of viral load measurements taken across all individuals. Abbreviations: “IN”: Intranasal; “CJ”: Conjunctival; “RM”: Rhesus Macaque; “CM”: Cynomolgus Macaque. *Using a conversion factor of 0.7, the approximate dose in units of TCID_50_ is 10^6.9^ TCID_50_.

| Study | Dose | Route | Strain | Species | Age | Sex | Swab Type(s) | No. of Pos. | No. of Obs. |
| --- | --- | --- | --- | --- | --- | --- | --- | --- | --- |
| Li et al. (2005) [11] | $10^5$ TCID<sub>50</sub> | IN | PUMC01 | RM | Juvenile | F, M | Oral | 8/8 (100%) | 8 |
| Lawler et al. (2006) [12] | $6 \times 10^6$ PFU* | IN+CJ | Urbani | CM | Adult | M | Nasal, Oral | 2/2 (100%) | 46 |
| Nagata et al. (2007) [13] | $10^3$ TCID<sub>50</sub> | IN | HKU-39849 | CM | Juvenile | M | Nasal, Oral | 0/2 (0%) | 16 |
| | $10^6$ TCID<sub>50</sub> | IN | HKU-39849 | CM | Juvenile | M | Nasal, Oral | 2/2 (100%) | 22 |
| Chen et al. (2008) [14] | $10^5$ TCID<sub>50</sub> | IN | PUMC01 | RM | Juvenile | F, M | Oral | 11/11 (100%) | 11 |
| Liu et al. (2016) [15] | $10^5$ TCID<sub>50</sub> | IN | PUMC01 | RM | Adult | F | Oral | 12/12 (100%) | 72 |
| Liu et al. (2019) [16] | $10^5$ TCID<sub>50</sub> | IN | PUMC01 | RM | Adult | F | Oral | 2/2 (100%) | 5 |

**Fig 1.**
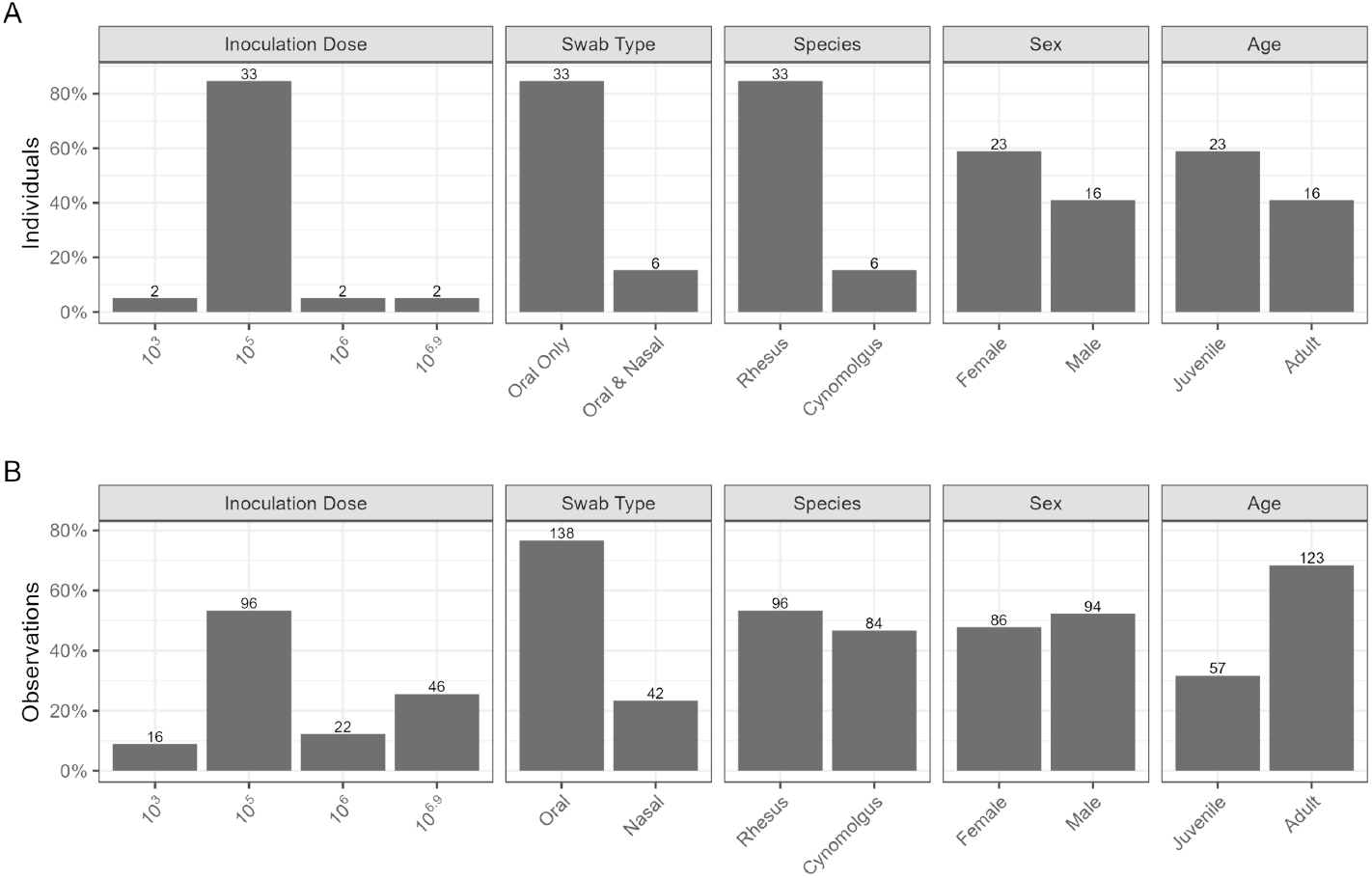
Summary of collected URT swab data from SARS-CoV-1 NHP challenge studies. Each panel shows the distribution of individuals (A) or observations (B) for a covariate in our dataset. The numbers over the bars indicate the number of individuals or observations in each covariate category. The inoculation doses are in units of TCID_50_.

To analyze these data, we developed a joint dose-response and kinetics model for SARS-CoV-1 infections in NHPs. Under this model, infection proceeds as a sequence of three steps: (i) an individual NHP is exposed to a dose *V*_0_ of SARS-CoV-1 (**Fig 2A**), (ii) the NHP is infected with probability *P* (*V*_0_) (**Fig 2B**), and (iii) if the NHP is infected, then virus grows and decays exponentially and can be detected via viral load values from URT swabs (**Fig 2C**). For the dose-response component of our model (**Fig 2B**), we used an exponential dose-response relationship to describe the probability of infection, which has a “hit probability” (per-virion successful infection probability) *k* as its key parameter. The ID_50_ can be computed from the hit probability via ID_50_ = ln(2)*/k*. For the viral kinetics component of our model (**Fig 2C**), we used a piecewise linear function on a logarithmic scale to approximate the kinetics; this approach has been previously used in other modeling studies [17–20]. We also allowed the initial observed viral load *N*_0_ (the effective viral load at 0 days post infection, representing the subset of the inoculum that establish the successful infection) and the time of peak observed viral load *t*_*p*_ to vary by dose. In particular, we used log-log linear equations to model the relationship between the expected number of successful virions at each dose level and the values of *N*_0_ and *t*_*p*_. Slope parameters *α*_*N*_ and *α*_*t*_ control the strength and direction of dose effects on *N*_0_ and *t*_*p*_, respectively. Positive values of the slope parameters indicate that *N*_0_ and *t*_*p*_ are positively related to exposure dose; negative values indicate a negative relationship.

**Fig 2.**
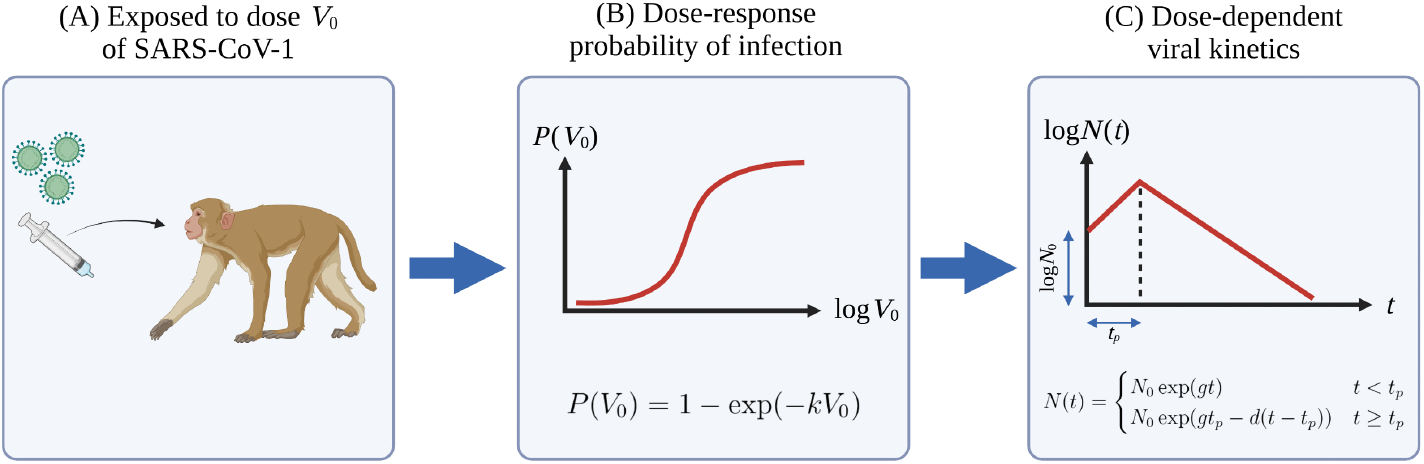
Dose-response and viral kinetics model for SARS-CoV-1 inoculations in non-human primates. An individual non-human primate is exposed to a dose *V*_0_ of SARS-CoV-1 (A), which results in a probability of infection given by the exponential dose-response curve (B). Given a successful infection, viral RNA can be observed in URT swabs as measured viral load values, which follow a pattern of exponential growth and decay (C).

To aggregate NHP viral load data across studies, we included study-level hierarchy for many of our model parameters. The hierarchical model structure provided flexibility to account for systematic lab and methodological differences between studies, and it also allowed us to assess overall dose effects on viral kinetics across all studies. Full model details are available in the **Methods** and **S1 Methods**. We used Bayesian inference via Markov Chain Monte Carlo to fit our model to the collected URT viral load data, which allowed us to jointly infer viral kinetics and dose-response parameters.

### Study-Specific Viral Kinetics Fits

The hierarchical structure of our model allowed us to simulate predicted viral load trajectories in URT swabs for each of the studies included in our analysis (**Fig 3A**). Overall, the posterior predicted viral load trajectories aligned reasonably well with the observed data. As expected, there was less uncertainty in the predicted trajectories for studies that measured more extended time series data. For example, the 10^6.9^ TCID_50_ dose individuals from Lawler et al. (2006) had many reported measurements at later time points, so the predicted viral load trajectories for these individuals clustered together closely even at later times. Conversely, among all of the studies that had 10^5^ TCID_50_ dose individuals, there was only one viral load measurement taken past seven days post infection, so there was more uncertainty in the exponential decay phase of the viral kinetics for these individuals. None of the 10^3^ TCID_50_ dose individuals in Nagata et al. (2007) had detectable viral RNA. The obvious interpretation is that these individuals were not infected, but the model allows for the possibility of successful infections that escaped detection. Predicted trajectories for these individuals (assuming successful infection) had to be extrapolated from higher-dose individual data, which led to greater uncertainty in these predictions. Notably, the predicted successful viral load trajectories for the 10^3^ TCID_50_ dose individuals in Nagata et al. (2007) were below the effective limit of detection (i.e., the smallest reported viral load value from Nagata et al. (2007)). Our model thus suggested that some 10^3^ TCID_50_ dose individuals from Nagata et al. (2007) could plausibly have been infected but had viral loads too low to be detected by the qPCR assays used.

**Fig 3.**
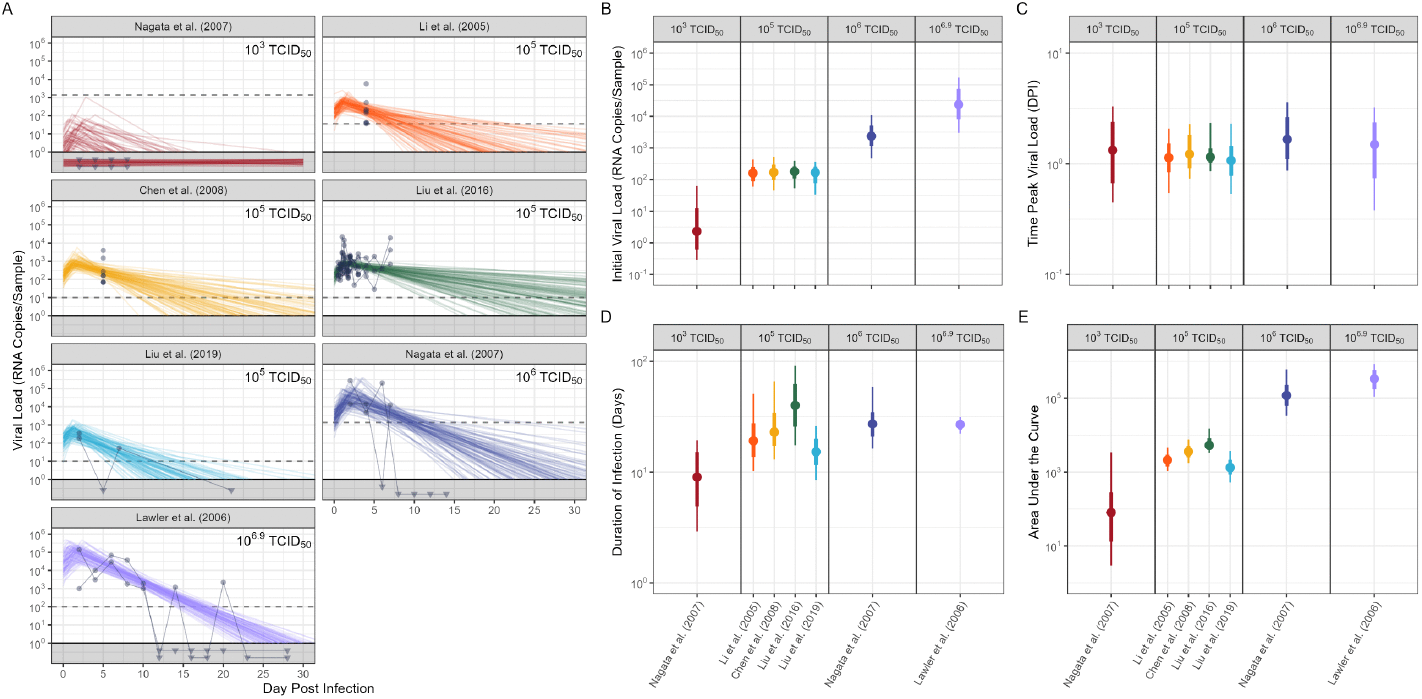
Results for study-specific predicted URT swab viral load trajectories. (A) Each panel shows the observed SARS-CoV-1 viral load trajectories in oral swabs for a study and dose combination. Each point represents a measured viral load value from an oral swab, and lines connecting multiple points represent measurements taken from the same individual over time. Upside down triangles plotted in the gray region below 10^0^ indicate measurements below the limit of detection (LOD). For the Lawler et al. (2006) panel, the horizontal dotted line indicates the reported LOD; for all other panels, the horizontal dotted lines indicate the effective LOD (smallest measured value from that study). Semitransparent colored lines are 100 random draws from the inferred within-host viral load kinetics (i.e., predicted viral load trajectories based on parameter sets drawn from the joint posterior distributions), and lines that fall in the gray region indicate predicted failed infections. See **S2 Fig** for individual plots of all observed data. (B), (C), (D), and (E) show distributions of various metrics of the simulated successful infection trajectories. These metrics are the initial viral load (the effective viral load at 0 days post infection), the time of peak viral load, the duration of infection (time spent with viral load of at least 1), and area under the curve. The points represent the posterior median values, and the thick and thin bars indicate the 66% and 95% credible intervals, respectively.

Individuals that received larger inoculation doses were predicted to have higher initial viral loads (**Fig 3B**) and greater cumulative viral load (as measured by area under the viral load curve; **Fig 3E**). On the other hand, there did not appear to be a strong dose effect on the time of peak infection, with significant overlaps in the peak time credible intervals for all simulated viral load trajectories (**Fig 3C**). The relationship between inoculation dose and duration of infection (i.e., total time spent with a viral load of at least 1) was less clear (**Fig 3D**). Due to the sparsity of data at later time points, it was difficult to estimate viral load decay rates for many of the studies, which resulted in wide credible interval estimates for the infection durations.

### Dose Effects on Viral Load Kinetics

In addition to providing study-level predictions, the hierarchical structure of our model allowed us to assess the overall effect of inoculation dose on the resulting viral load trajectories across all studies. Our model predicted a clear positive relationship between the initial viral load and inoculation dose (**Fig 4A**), such that larger inoculation doses led to larger initial viral loads. The median slope parameter for the initial viral load (*α*_*N*_) on this log-log plot was 1.09 (95% CrI: [0.59, 1.56]). The fact that the median value for *α*_*N*_ was roughly one indicated that an approximately linear relationship between the inoculation dose and initial viral load was a highly plausible explanation for the patterns in our dataset. For example, the median fold change in initial viral load after a 100-fold increase in inoculation dose (from 10^5^ TCID_50_ to 10^7^ TCID_50_) was 151.32 (95% CrI: [15.25, 1299.87]; **Fig 4C**).

**Fig 4.**
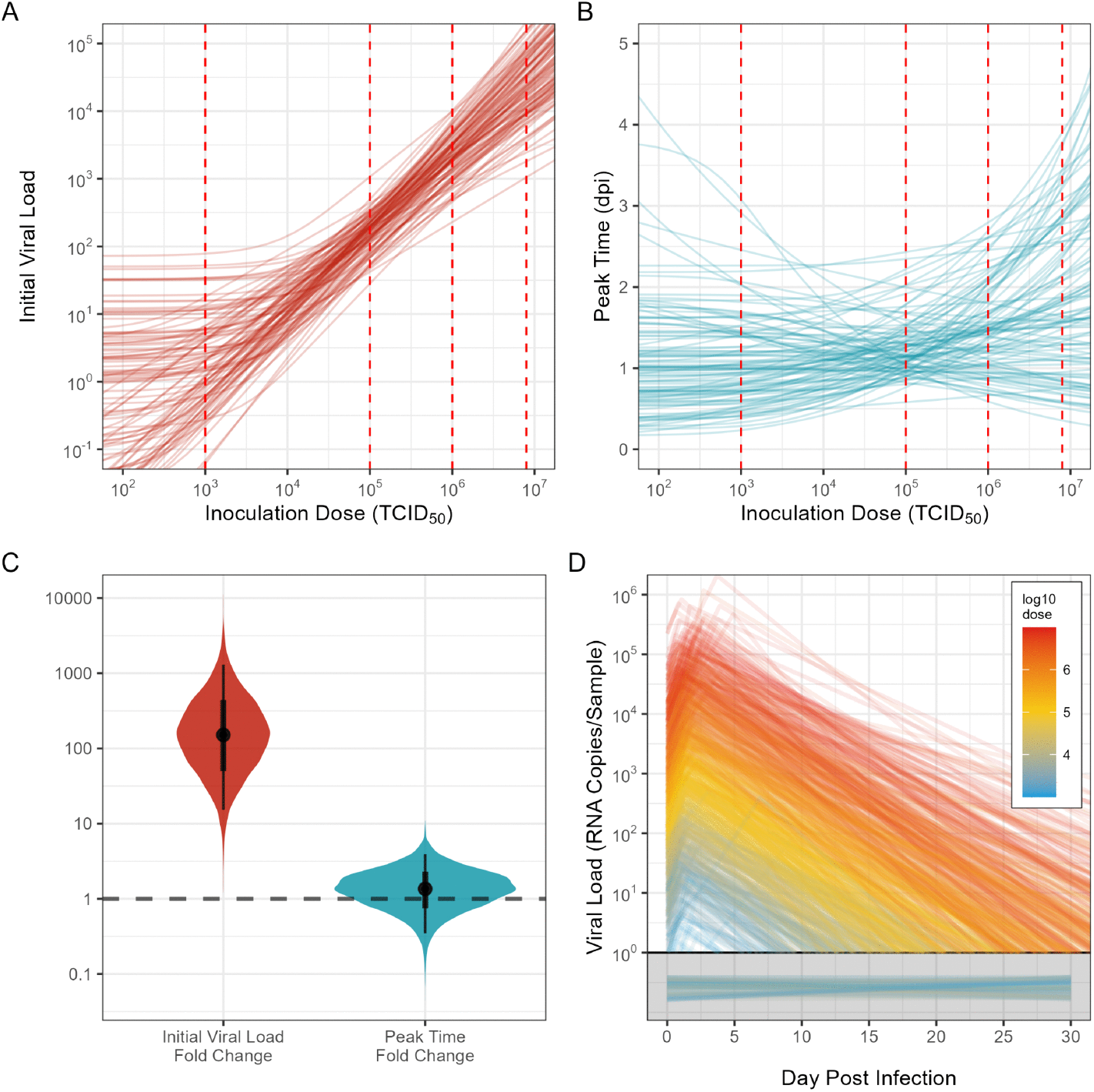
Results for effects of dose on viral kinetics. Relationship between the initial viral load (A) and time of peak viral load (B) with the inoculation dose. Semitransparent lines are 100 random draws from the inferred posterior relationship. The non-linearity of these inferred curves comes from the fact that we encoded the log initial viral load and log peak time as functions of the mean of the zero-truncated Poisson distribution of the number of successful virions (i.e., these plots were generated assuming that infection was successful, see **Methods** for additional details). Dotted red vertical lines indicate doses that were used in the selected non-human primate challenge studies. (C) Inferred distributions of the effect of a 100-fold dose change (10^5^ to 10^7^ TCID_50_) on the initial viral load and time of peak viral load. The dotted gray horizontal line indicates no effect of inoculation dose on the viral kinetics. (D) Random draws from the inferred model average (non-study-specific) within-host viral load kinetics (i.e., predicted viral load trajectories based on parameter sets drawn from the joint posterior distributions). We plotted 1,000 random draws with inoculation doses ranging from 10^3^ to 10^7^ TCID_50_. Lines that fall in the gray region below 10^0^ indicate predicted failed infections.

Conversely, our model did not predict a clear relationship between the timing of peak viral load and inoculation dose (**Fig 4B**). The median slope parameter for the timing of peak viral load (*α*_*t*_) was 0.07 (95% CrI: [−0.23, 0.30]). The median fold change in peak timing after a 100-fold increase in inoculation dose (from 10^5^ TCID_50_ to 10^7^ TCID_50_) was 1.36 (95% CrI: [0.35, 3.93]; **Fig 4C**). A highly plausible explanation for the patterns of viral load kinetics in our dataset was that the inoculation dose did not impact the timing of peak viral load. However, the alternatives where peak timing was either positively or negatively associated with inoculation dose cannot be ruled out. When we fit our model using less-informative priors, these slope parameters had wider posterior distributions, but our conclusions remained unchanged (**S3 Fig**).

The combined effects of inoculation dose on initial viral load and time of peak viral load can be visualized by simulating average (non-study-specific) viral load trajectories (**Fig 4D**). As the inoculation dose increased, the simulated viral load trajectories tended to start at larger initial viral load values, and the time of peak viral load was similar across inoculation doses.

### Estimates of Infectious Dose

From the dose-response component of our model, we estimated a posterior median ID_50_ of 10^3.03^ TCID_50_ (95% CrI: [10^0.30^, 10^4.24^]; **Fig 5A**). Almost all of the individuals in our compiled data were inoculated with doses of at least 10^5^ TCID_50_ (37 out of 39 NHPs), and all of these individuals had detectable viral RNA following inoculation (**Table 1**), which provided strong information that these individuals were all successfully infected. Thus, the majority of the ID_50_ posterior probability mass was below 10^5^ TCID_50_. Due to the small number of low-inoculation dose individuals, a range of ID_50_ values below 10^3^ TCID_50_ were plausible. Additional experiments at lower dose ranges would allow us to further refine the lower bound on the ID_50_ estimates.

**Fig 5.**
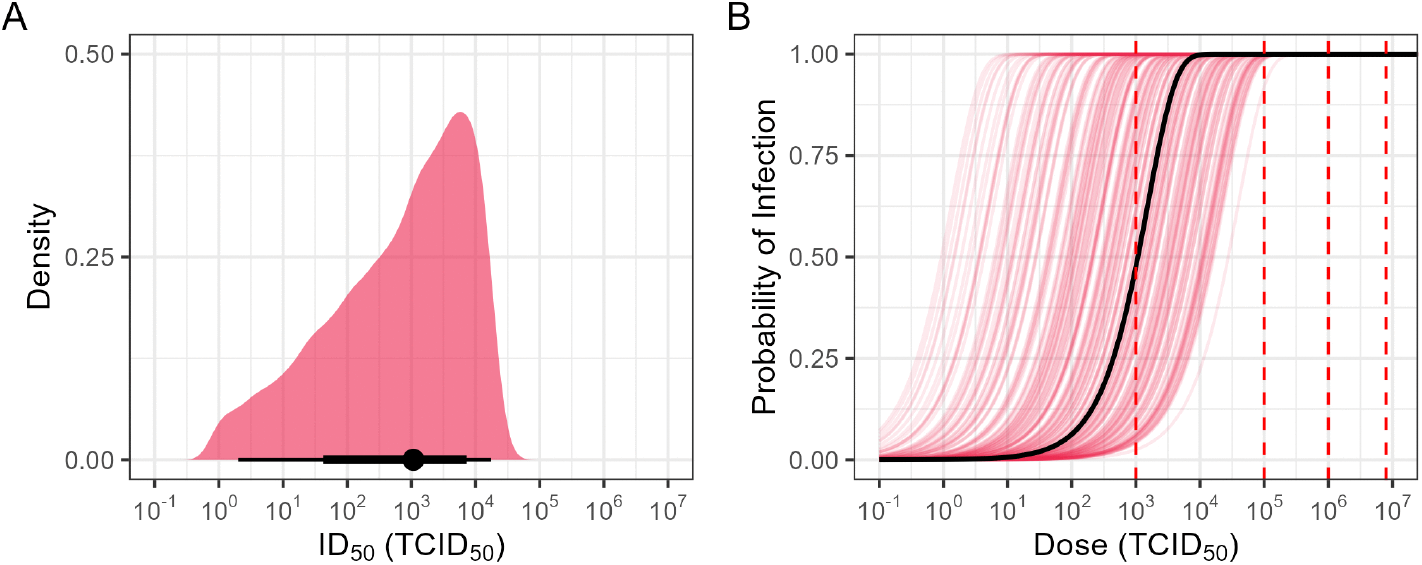
Estimated infectious doses and infection probabilities. (A) Inferred posterior distribution of infectious dose 50 (ID_50_) values for SARS-CoV-1 infections in non-human primates following intranasal inoculations. The black point represents the median ID_50_ value, and the thick and thin black bars indicate the 66% and 95% credible intervals, respectively. (B) Exponential dose-response curves for 200 sampled hit probabilities from the inferred posterior distribution. The black curve represents the median estimated dose-response curve. The dotted red vertical lines indicate doses used in the included studies.

Using the estimated dose-response curves, the four inoculation doses represented in the data (10^3^, 10^5^, 10^6^, 10^6.9^ TCID_50_) had posterior median probabilities of infection of 0.48 (95% CrI: [0.04, 1.00]), 1.00 (95% CrI: [0.98, 1.00]), 1.00 (95% CrI: [1.00, 1.00]), and 1.00 (95% CrI: [1.00, 1.00]), respectively. For individuals inoculated with doses greater than 10^5^ TCID_50_, our model predicted that successful infection was essentially guaranteed, but for lower doses there was greater uncertainty in the estimate for infection probability, with a much wider distribution of estimated values (**Fig 5B**). The wider distribution of estimated probabilities for low-inoculation dose individuals was another consequence of the wide range of plausible low ID_50_ values in the sampled posterior draws, which in turn resulted from the sparse data at doses below 10^5^ TCID_50_.

### Sensitivity to Variations in Study Design

We repeated our model-fitting procedure with various subsets of the combined dataset. This allowed us to assess what information was contributed by different studies, and it also allowed us to test for potential sources of bias.

We first subsetted our data by swab type (138/180 observations for oral swabs, and 42/180 observations for nasal swabs). When we fit our model to only oral swab data, we obtained a median ID_50_ of 10^2.99^ TCID_50_ (95% CrI: [10^0.37^,10^4.23^]; **Fig 6A**), which was very similar to the estimates obtained using the full dataset. When we fit our model to only nasal swab data, we obtained a higher median ID_50_ of 10^4.15^ TCID_50_ (95% CrI: [10^1.03^, 10^5.79^]; **Fig 6A**). This increase was expected as the nasal swab subset did not include any 10^5^ TCID_50_ dose individuals. All of the 10^5^ TCID_50_ dose individuals had detectable viral RNA in their oral swabs and were therefore very likely successfully infected, providing strong evidence that the ID_50_ is well below 10^5^ TCID_50_. Without this data, the model had trouble rejecting larger ID_50_ values.

**Fig 6.**
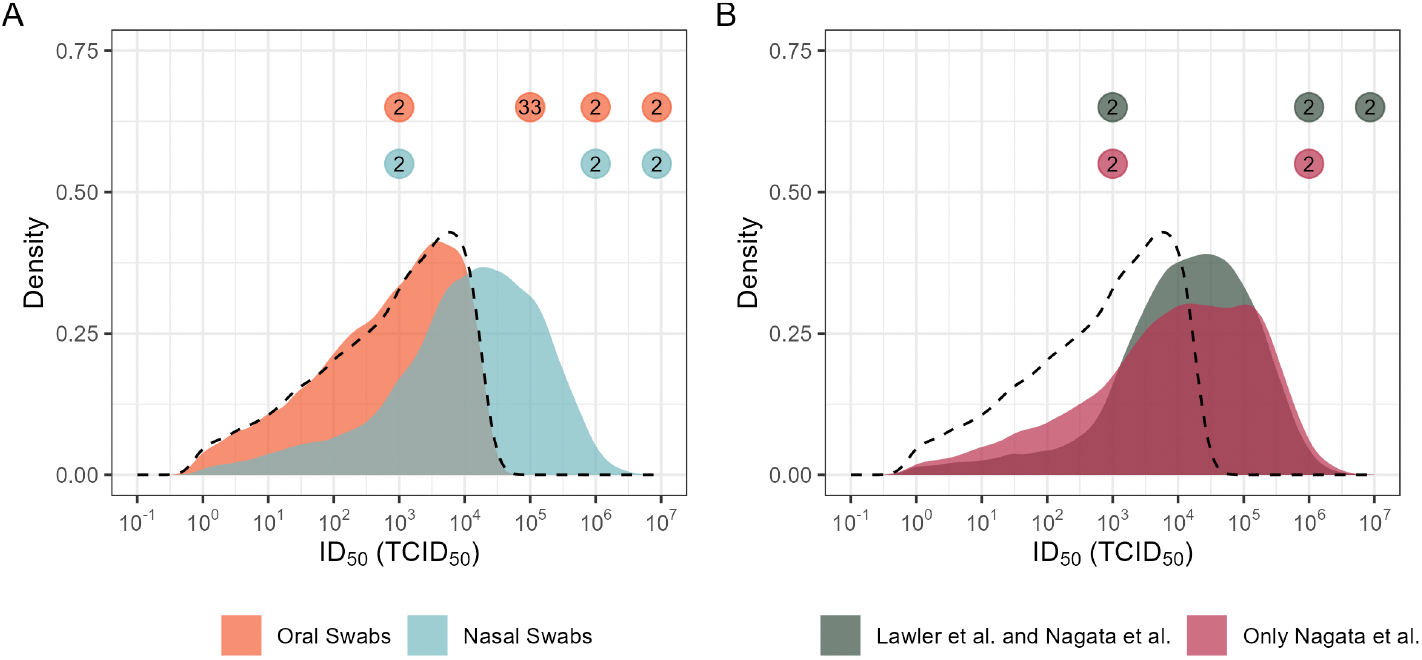
Results of infectious dose sensitivity analysis. (A) Inferred posterior distributions of infectious dose 50 (ID_50_) values using data from different swab types in our dataset. (B) Inferred posterior distributions of ID_50_ values using data from only cynomolgus macaques (data from Lawler et al. (2006) and Nagata et al. (2007)) and data from only Nagata et al. (2007). The colored circles above the densities indicate the number of individual NHPs per dose level in each of the data subsets. For reference, the dashed black line indicates the posterior ID_50_ density when the entire dataset is used (**Fig 5A**).

Next, we subsetted our data by NHP species. The cynomolgus macaque observations in our dataset (84/180 observations) came from two different studies (Lawler et al. (2006) and Nagata et al. (2007)) and included three different doses (10^3^, 10^6^, 10^6.9^ TCID_50_). Similar to when we fit our model to nasal swab data only, since none of the cynomolgus macaques were exposed to an inoculation dose of 10^5^ TCID_50_, our model inferred a larger posterior median ID_50_ of 10^4.19^ TCID_50_ (95% CrI: [10^1.09^, 10^5.75^]; **Fig 6B**). We also fit our model using only data from Nagata et al. (2007), since this was the only study included in our analysis where NHPs were inoculated intranasally with more than one inoculation dose. The median ID_50_ in this case was 10^4.06^ TCID_50_ (95% CrI: [10^0.77^, 10^5.84^]; **Fig 6B**). All rhesus macaques in our dataset were exposed to the same inoculation dose (10^5^ TCID_50_), so we did not fit our model to observations from rhesus macaques only.

Estimates of the other parameters were broadly consistent across the different studies and data subsets. The growth rate estimates were similar for all studies, regardless of the data subset (**S5 Fig**). The decay rate estimates were more varied across the studies, although the 95% credible intervals still overlapped across all studies and subsets (**S5 Fig**). This increased variability was expected since there were very few observations collected at later time points, which limited the precision with which decay rates could be estimated. The slope parameters governing the dependence between the inoculation dose and viral kinetics were also broadly similar across data subsets (**S6 Fig**); the median values for *α*_*N*_ were all around one, and the 95% credible intervals for *α*_*t*_ overlapped zero in all data subsets.

Two additional key parameters are *β*_*N*_ and *β*_*t*_, which represent the log initial viral load and log time of peak viral load, respectively, given an inoculation dose of 10^5^ TCID_50_ (see **Methods** for additional details). There were three data subsets (only nasal swab data; only cynomolgus macaque data; and only Nagata et al. (2007) data) that did not include any 10^5^ TCID_50_ dose individuals. For these subsets, the 95% credible intervals for *β*_*N*_ and *β*_*t*_ were wider than those obtained using the full dataset (**S7 Fig**), which reflected the need to extrapolate these parameters from other dose levels. Importantly, the 95% credible intervals estimated using the full dataset substantially overlapped with (and were often entirely contained within) the wider intervals obtained from subsets lacking the 10^5^ TCID_50_ dose individuals. This indicated that although uncertainty increased when specific dose groups were excluded, no individual study or subset appeared to systematically bias the parameter estimates.

## Discussion

In this study, we developed a joint dose-response and viral kinetics modeling framework to analyze data from SARS-CoV-1 NHP challenge studies. Our model included study-level hierarchy for several key parameters, which allowed us to aggregate NHP viral load data across multiple studies in order to obtain a sufficient number of individuals and range of inoculation doses to perform a dose-response analysis. Our model also allowed us to investigate potential dose effects on viral kinetics.

We obtained a median ID_50_ estimate of 10^3.03^ TCID_50_ for SARS-CoV-1 via intranasal inoculations in rhesus and cynomolgus macaques. Our estimate of the ID_50_ is the first-ever estimate for the infectious dose of SARS-CoV-1 in NHPs. Previously, a dose-response model for SARS-CoV-1 was developed by Watanabe et al. (2010), where they estimated the infectious dose of SARS-CoV-1 in transgenic mice [4]. In their model, they used an exponential dose-response curve fit to a combined dataset from transgenic mice (expressing human ACE2 in all tissues) challenged with SARS-CoV-1 and unaltered mice challenged with the mouse virus MHV-1. They estimated an ID_50_ of 280 PFU (using death as the endpoint), which was much lower than our median estimate. The transgenic mice used in this study have been shown to be highly susceptible to SARS-CoV-1, with infectious virus being detected in tissues not associated with natural infections. For example, virus was detected in the brains of the mice following inoculation [21, 22]. Given the greater relatedness and physiological similarities between NHPs and humans, NHP challenge data are likely to provide a better proxy for respiratory infectiousness of SARS-CoV-1 in humans [23–25].

For the URT viral kinetics, our model predicted that larger inoculation doses resulted in higher initial viral loads, which meant that infections resulting from larger exposure doses tended to start with larger concentrations of detectable viral RNA. However, our model was not able to detect an effect of inoculation dose on the timing of peak viral load (i.e., the time when the concentration of viral RNA is at a maximum). Previous experimental challenge studies have examined the relationship between inoculation dose and viral loads, with findings that vary by animal species and pathogen. For example, inoculation dose and viral loads were associated in mouse models of respiratory viruses [26–28], as well as in rhesus macaque studies of Zika virus [29] and human challenge studies of norovirus [30]. On the other hand, studies in African green monkeys found no association between dose and viral loads for SARS-CoV-2 [31], and studies in hamsters found no association for both SARS-CoV-1 [32] and SARS-CoV-2 [33].

One of the main challenges of this analysis was developing a principled way to aggregate and utilize data across the different studies. The most common data type collected and reported in the NHP SARS-CoV-1 challenge studies was viral load data, as determined by quantitative PCR, so we used these data in our analysis. These data are expected to be well correlated with presence of infectious virus [34], though there is evidence that the dose-response curves can be quite different depending on whether PCR or culture results are used [9]. Incorporating additional data types (e.g., serology, infectious virus titers) could yield more precise parameter estimates. For example, there were very few low-dose individuals in our dataset, so our model had difficulty refining the lower bound for the ID_50_ estimates. The low-dose individuals were all from Nagata et al. (2007). The authors reported that those individuals did not seroconvert, had no detectable infectious virus or viral RNA (nasal, oral, rectal swabs) up to 8 dpi, and had no detectable antibodies (by indirect fluorescence and neutralizing antibody tests) up to 8 dpi. Thus, they concluded that infection failed to establish in these individuals [13]. This type of information could potentially be incorporated into future versions of our model (i.e., the likelihood computation) and could help with tightening the ID_50_ estimates by increasing the probability that the low-dose individuals were non-infected. Since these additional measurements were not conducted or reported in the other studies in our dataset, we did not use them in this first analysis.

Another limitation of the NHP data was that individual-level time series data was rare. Out of the 39 individuals in our combined dataset, 20 individuals had only a single viral load measurement, and only 10 of the individuals had viral load measurements taken on or after 7 dpi. The lack of extensive time series data, especially measurements for later times, made it challenging to estimate viral load decay rates and the overall viral load trajectories. Additionally, the lack of data resolution meant that we could not definitively determine whether there was or was not any dose dependence on the timing of the viral kinetics, and we ultimately relied on a relatively crude model for the viral kinetics (piecewise linear on a logarithmic scale). With a richer dataset with more time series data, our model could be extended to capture individual-level heterogeneity in viral kinetics. The two individuals from Lawler et al. (2006) had the most time points (11 measurements taken from 2 to 28 dpi). Those data in particular show evidence of individual-level heterogeneity; one of the individual’s infection peaked substantially earlier than the other’s. A richer set of time series data could allow us to quantify the degree of heterogeneity across individuals, which represent differential responses to infection and are key to understanding phenomena such as superspreading [35].

We sought to minimize the influence of study-level differences by using individuals from studies with similar experimental procedures (similar species, PCR procedures, etc.), and by developing a Bayesian hierarchical model to capture and account for any systematic differences among studies. We repeated our model-fitting procedure with several subsets of our full combined dataset, and the estimates for key parameters were similar across all studies regardless of the data subset. However, we cannot rule out the possibility that unwanted strain/species/lab differences contributed to the patterns in our results. There is some evidence from previous studies that cynomolgus macaques experience more severe infections than rhesus macaques following SARS-CoV-1 inoculation [36, 37], but small sample sizes and differences in viral strains in those studies prevented any definitive conclusions. In our case, fitting the dose-response and viral kinetics model to only cynomolgus macaque data resulted in similar parameter estimates; credible intervals were wider than when data from both cynomolgus and rhesus macaques were used but is explained directly by the distribution of doses tested. For the different SARS-CoV-1 virus strains, it was difficult for us to assess whether there were any strain effects since the only studies that used different strains also used different macaque species. The inferred posterior distributions for growth and decay rates across the studies and strains in our analysis were all similar (**S5 Fig**), which suggested that the virus strain differences were minor. A previous study in hamsters found that intranasal inoculation with the Urbani and HKU-39849 strains resulted in similar viral titers, and these two strains have identical spike protein amino acid sequences [38]. Additionally, for studies in our analysis that reported details on RT-PCR targets for viral load measurements, we verified that all primers targeted the same orf1ab region (verified using SnapGene software; www.snapgene.com). However, due to limitations in the available data, there is the possibility that our model was unable to distinguish the finer differences in kinetics between the various SARS-CoV-1 strains.

Overall, our study proposes a new approach to studying dose-response relationships and viral kinetics even when data are sparse and fragmented across multiple studies, and it presents the first ever dose-response analysis of SARS-CoV-1 in NHPs. Despite some limitations in the available NHP SARS-CoV-1 data, we were able to produce useful bounds on the ID_50_ in NHPs and identify an effect of inoculation dose on initial viral load. The joint dose-response and viral kinetics modeling framework we developed is extremely flexible and could be expanded in various ways. In the current model, we have only used viral load data from URT swabs to inform the likelihood, but it would be straightforward to incorporate additional sources and types of data to compute the probabilities of infection. The hierarchical aspect of our modeling approach allowed us to reuse virological data from multiple previous challenge studies and draw additional insight from the valuable quantitative measurements collected in those experiments. The approach can also be applied to single studies that measure outcomes from multiple dose levels to make better use of the rich data collected and achieve greater statistical power than standard analyses that treat infection as a binary outcome. This may be particularly valuable in instances where it is unclear how to classify individuals as infected or non-infected. Our modeling framework can be adapted readily to other host-pathogen systems and contributes to a growing movement toward mechanistic analysis of dose-response relationships [1], with the aim of maximizing the scientific value derived from animal experiments and hence reducing the demand for further experiments in line with 3Rs principles [39].

## Methods

### Data Collection

We performed a comprehensive literature search for NHP challenge studies and identified 37 studies (see **S1 Methods** for further details). To be included in the analysis herein, we selected articles from the broader dataset that met the following criteria:

1. Be a primary study involving experimental infection of rhesus macaques (*Macaca mulatta*) or cynomolgus macaques (*Macaca fascicularis*) with a strain of SARS-CoV-1 that had not been genetically modified.
2. Report qualitative and/or quantitative viral load data from URT swabs.

In total, six studies were included in our dose-response and viral kinetics analysis (**Table 1**).

Raw data was not published in any of the six included studies, so we manually extracted viral load values from the texts, tables, and figures. We extracted data from figures using the R package digitize (version 0.0.4) [40]. All studies reported inoculation doses in TCID_50_ except for Lawler et al. (2006), which used units of PFU. For that study, we used a conversion factor of 0.7 to convert the reported PFU dose to an approximate TCID_50_ value: 6 × 10^6^ PFU */*0.7 ≈ 10^6.9^ TCID_50_ [41]. We recorded additional covariates for each individual NHP, including age and sex. For the NHP age groups, “juvenile” included individuals with reported ages between 2 and 4 years, and “adult” included individuals reported as adults but without numeric ages.

### Joint Dose-Response and Kinetics Model

We developed a model to simultaneously perform a dose-response analysis and a within-host kinetics analysis of SARS-CoV-1 in NHPs.

We used a single-hit, independent action, exponential dose-response relationship for the SARS-CoV-1 infection probability in NHPs following intranasal inoculation. Under this model, if the host is exposed to *V*_0_ virions, then the number of successful virions, *V*_*s*_, is Poisson distributed with mean *kV*_0_, where *k* is the hit probability (i.e., the per-virion success probability) [1]. The probability that the host is infected (*V*_*s*_ *>* 0) after exposure to *V*_0_ virions is

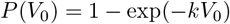

Given a successful infection, we modeled within-host viral dynamics as exponential growth up to a peak time and then exponential decay afterwards. The mean measured viral load from URT swabs at day *t* is

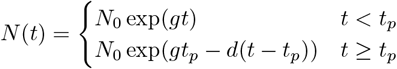

where *N*_0_ is the mean initial viral load established immediately following inoculation (i.e., the effective viral load at 0 dpi), *t*_*p*_ is the time of peak viral load, *g* is the growth rate, and *d* is the decay rate. Observed viral load measurements were assumed to be log-normally distributed around the mean viral loads:

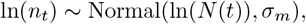

where *n*_*t*_ is an observed viral load measurement at time *t*, and the standard deviation *σ*_*m*_ accounts for measurement variation. For unsuccessful infections, we assumed a constant false positive rate of detecting viral RNA over all days.

We also included the possibility that the initial viral load (*N*_0_) and time of peak viral load (*t*_*p*_) were dependent on the inoculation dose. First, given that infection was successful (*V*_*s*_ *>* 0), the expected number of successful virions is

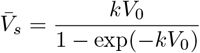

which is the mean of the zero-truncated Poisson distribution of *V*_*s*_. Next, we encoded the dose-dependence of the initial viral load and time of peak viral load with log-log linear relationships:

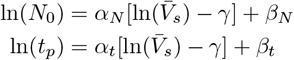

The magnitude and sign of the slope parameters (*α*_*N*_ and *α*_*t*_) control the strength and direction of the dose-dependence of the initial viral load and time of peak viral load. To improve the interpretability of the intercept parameters (*β*_*N*_ and *β*_*t*_) we included a horizontal shift of *γ*, which is the log expected number of successful virions (given a successful infection) following an inoculation dose of 10^5^ TCID_50_, i.e., 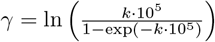. The intercept parameters thus represent the log initial viral load and log time of peak viral load given an inoculation dose of 10^5^ TCID_50_. By encoding the dependence in this way, we allowed for several possible relationships between the viral kinetics parameters and the initial inoculation dose.

### Accounting for Between-Study Differences

To account for lab and methodological differences across studies, we included study-level hierarchy for many of our model parameters. Not all studies used the same SARS-CoV-1 strains and macaque species, so we allowed the growth and decay rate parameters to be different for each study. We also allowed the *β*_*N*_ and *β*_*t*_ intercept parameters in the viral kinetics dose-dependence equations to vary between studies. Finally, we allowed the magnitude of variation in observed measurements (Gaussian noise) to be study-specific.

Although this hierarchical structure is capable of capturing systematic differences across studies, it is important to note that potential differences due to virus strain and host species will be jointly captured in the study-level variation parameters, so our approach cannot readily separate out strain and species effects.

### Computational Methods

Data preparation, analysis, and visualizations were completed with R version 4.3.3 [42]. The following packages were used for processing data and plotting results: dplyr, ggplot2, and tidybayes [43–45].

We used Markov Chain Monte Carlo methods to infer parameters in our joint dose-response and viral kinetics model. We implemented and conducted inference using the Stan platform [46], and posterior samples were drawn via the R interface CmdStanR version 0.7.0 [47]. We used four Markov chains, each with 5000 iterations. The first 1000 iterations were used as the warmup period, and the last 4000 iterations were used for parameter estimation. We assessed model convergence using 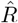 and effective sample sizes. We also inspected mixing of chain traces visually. No issues were detected. Detailed descriptions of the likelihood calculation and prior selections are available in the **S1 Methods**.

## Data and Code Availability

All data and code used to produce the results and figures in this manuscript are available on GitHub at: https://github.com/pcl1701/sars1-dose-response.

## Acknowledgments

J.O.L.-S., C.E.S and D.H.M. were supported by the Defense Advanced Research Projects Agency DARPA PREEMPT #D18AC00031. C.E.S was also supported by the National Institutes of Health (grant 5T32 GM008185-33) and the UCLA Office of the Vice Chancellor for Research 3R Grant. J.O.L.-S. was also supported by the National Science Foundation (DEB-2245631) and the UCLA AIDS Institute and Charity Treks. The funders had no role in study design, data collection and analysis, decision to publish, or preparation of the manuscript. The content of the article does not necessarily reflect the position or the policy of the US government, and no official endorsement should be inferred.

## Competing Interests

J.O.L.-S., P.C.L., and C.E.S. declare no competing interests. D.H.M. is an employee of the United States Centers for Disease Control and Prevention (CDC), but conducted this work in a personal capacity, as an independent consulting scientist working with the J.O.L.-S. lab at UCLA. The findings and conclusions in this manuscript do not necessarily represent the scientific positions of the CDC or the United States government.

## Supporting information

**S1 Fig. Screening and selection procedure for SARS-CoV-1 non-human primate challenge studies**. Studies were processed following the Preferred Reporting Items for Systematic Review and Meta-Analyses (PRISMA) guidelines.

**S2 Fig. Observed viral load trajectories for all individuals**. Each panel displays the reported viral load data for a single non-human primate (NHP) included in our analysis. Each individual NHP has a unique label composed of the acronyms of the first author (from the source study), a species acronym, and a number. The dashed horizontal lines indicate the limit of detection (for Lawler et al. (2006)) or the effective limit of detection (for all other studies). Upside down triangles plotted in the gray region indicate measurements below the limit of detection. Data for three individuals from Chen et al. (2008) are not displayed as those individuals each only had a single “detected” (non-quantitative) measurement. Study acronyms: “BL”: Li et al. (2005); “JL”: Lawler et al. (2006); “LL”: Liu et al. (2016), Liu et al. (2019); “NN”: Nagata et al. (2007); “YC”: Chen et al. (2008). Species acronyms: “RM”: rhesus macaque; “CM”: cynomolgus macaque.

**S3 Fig. Prior sensitivity analysis for dose-dependence parameters**. Prior sensitivity analysis for (A) the initial viral load slope parameter (*α*_*N*_) and (B) the peak viral load time slope parameter (*α*_*t*_). We tested the effect of using priors of increasing width on the final posterior estimates (setting different priors for the 10^7^:10^5^ dose-effect ratio). The left-hand side of each violin is the prior, and the right-hand side of each violin is the posterior. The first column (“1x SD”) is the original prior used to generate the main text results. The second column is for a prior of two times the standard deviation of the original prior, and the third column is for a prior of three times the standard deviation of the original prior.

**S4 Fig. Prior and posterior checks for SARS-CoV-1 dose-response and viral kinetics model**. Prior (A) and posterior (B) simulated mean viral load trajectories for each study and dose combination; semitransparent blue lines are 100 random draws from the prior/posterior distributions, and lines that fall in the gray region indicate failed infections. Prior (C) and posterior (D) simulated viral load measurements for each study and dose combination; semitransparent red points are observed viral load measurements taken from oral swabs (upside down triangles plotted in the gray region indicate measurements below the limit of detection), and red lines connecting multiple points represent measurements taken from the same individual over time; semitransparent blue points connected by blue lines are 100 simulated observed measurements from the prior/posterior distributions, and points/lines that fall in the gray region indicate failed infections. The dashed horizontal lines indicate the limit of detection (for Lawler et al. (2006)) or the effective limit of detection (for all other studies). Study acronyms: “BL”: Li et al. (2005); “JL”: Lawler et al. (2006); “LL16”: Liu et al. (2016); “LL19”: Liu et al. (2019); “NN”: Nagata et al. (2007); “YC”: Chen et al. (2008).

**S5 Fig. Posterior draws of growth/decay rates and doubling/halving times for different subsets of data**. Inferred posterior distributions of (A) growth rates, (B) decay rates, (C) doubling times, and (D) halving times for model fits on different subsets of our non-human primate dataset. These parameters have study-level hierarchy in our viral kinetics model, so each fit contains up to six inferred posteriors for the parameters (one posterior distribution per study). Points indicate the median posterior estimates. Thick and thin bars indicate the 66% and 95% credible intervals, respectively. Study acronyms: “BL”: Li et al. (2005); “JL”: Lawler et al. (2006); “LL16”: Liu et al. (2016); “LL19”: Liu et al. (2019); “NN”: Nagata et al. (2007); “YC”: Chen et al. (2008).

**S6 Fig. Posterior draws of dose-dependence slope parameters for different subsets of data**. Inferred posterior distributions of (A) initial viral load slope parameters (*α*_*N*_) and (B) peak viral load time slope parameters (*α*_*t*_) for model fits on different subsets of our non-human primate dataset. Points indicate the median posterior estimates. Thick and thin bars indicate the 66% and 95% credible intervals, respectively.

**S7 Fig. Posterior draws of dose-dependence intercept parameters for different subsets of data**. Inferred posterior distributions of (A) initial viral load intercept parameters (*β*_*N*_) and (B) peak viral load time intercept parameters (*β*_*t*_) for model fits on different subsets of our non-human primate dataset. These parameters have study-level hierarchy in our viral kinetics model, so each fit contains up to six inferred posteriors for the parameters (one posterior distribution per study). Points indicate the median posterior estimates. Thick and thin bars indicate the 66% and 95% credible intervals, respectively. Study acronyms: “BL”: Li et al. (2005); “JL”: Lawler et al. (2006); “LL16”: Liu et al. (2016); “LL19”: Liu et al. (2019); “NN”: Nagata et al. (2007); “YC”: Chen et al. (2008).

**S1 Methods. Additional methodological details, including literature search, likelihood computation, and prior justifications**.

## Notes

### Summary of Updates

Formatting changes to main manuscript; supplemental files added.

