## Supplemental methods for "Data aggregation and mechanistic modeling enable dose-response analysis of SARS-CoV-1 in non-human primates"

### 1 S1 Methods

#### 1.1 Comprehensive Literature Search

We conducted a comprehensive literature search on July 14th, 2021 to identify experimental challenge studies involving non-human primates (NHPs). Note that the original scope of our search included studies for both SARS-CoV-1 and MERS-CoV; however, for the present analysis, the identified articles were subsequently filtered to include only those in which NHPs were inoculated with SARS-CoV-1. We followed the Preferred Reporting Items for Systematic Review and Meta-Analyses (PRISMA) guidelines [1].

Studies were collected from Web of Science (Core Collection) and PubMed. Searches were conducted using the following search string: *(SARS-CoV OR MERS-CoV OR SARS OR MERS) AND (primate* OR macaque* OR monkey* OR "macaca" OR "chlorocebus") NOT (SARS-CoV-2 OR COVID-19).* To correspond with the emergence of SARS-CoV-1 in 2002 [2], the search results from Web of Science were additionally restricted by date from January 1st, 2002 to July 14th, 2021, and the search results from PubMed were restricted by year from 2002 to 2021. The preprint servers bioRxiv and medRxiv were also queried on July 14th, 2021 for SARS-CoV and MERS-CoV NHP challenge studies. Two search strings were used for these platforms. The first search string was: *(SARS-CoV OR MERS-CoV OR SARS OR MERS) AND (primate* OR macaque* OR monkey*) NOT (SARS-CoV-2 OR COVID-19).* The second search string was: *(SARS-CoV OR MERS-CoV OR SARS OR MERS) AND ("macaca" OR "chlorocebus") NOT (SARS-CoV-2 OR COVID-19).* These search results were also restricted by date from January 1st, 2002 to July 14th, 2021.

In total, 1598 articles were identified through Web of Science, PubMed, and the preprint servers, and an additional 17 articles were identified through Google Scholar. After removing duplicates we were left with 1351 unique records. From these records we removed articles that did not generate any primary data (reviews, opinion pieces, and news articles). We also removed articles that were irrelevant based on the following inclusion criteria: (i) must involve experimental infection of rhesus macaques (*Macaca mulatta*), cynomolgus macaques (*Macaca fascicularis*), or African green monkeys (*Chlorocebus sabaeus*) with a strain of SARS-CoV-1 or MERS-CoV that had not been genetically modified, and (ii) must report quantitative and/or qualitative data from at least one assay that measures viral load (RT-PCR) or viral titers (plaque assays or endpoint titration) in biological samples collected from at least one individual receiving a control treatment (e.g., no treatment, saline). After the screening procedure, the final collection contained a total of 37 articles (26 SARS-CoV-1 studies, 11 MERS-CoV studies).

For the SARS-CoV-1 dose-response and viral kinetics analysis, we further restricted articles by requiring experimental infection via intranasal inoculation with a strain of SARS-CoV-1 that had not been genetically modified. We also limited the articles to those that reported qualitative and/or quantitative viral load (RT-PCR) measurements from nasal and/or oral swabs. In total, six studies were included in our dose-response and viral kinetics analysis (**S1 Fig**).

#### 1.2 Likelihood Computation for NHP Viral Load Data

Our dataset contained 180 total measurements, and these measurements included both numeric viral load values (which were always greater than 0) as well as qualitative binary values (i.e., detected or not detected, "+" or "−"). Suppose an individual NHP $i$ had viral load measurements $n_{i,t}$ at several times $t \in T_i$. There were two cases to consider for the data likelihood: (i) the viral load measurements were observed given the individual was infected and (ii) the viral load measurements were observed given the individual was non-infected.

##### 1.2.1 Likelihood for Successful Infections

For a given set of parameters, our model predicts mean viral load trajectories for each individual in our dataset (see main text **Methods** for kinetics model details), where each individual $i$ has predicted mean viral load values $N_i(t)$. Given that individual $i$ was successfully infected, we modeled numeric values of $n_{i,t}$ around our predicted mean values (i.e., values from the viral kinetics model) with log-normal distributions:

$$\ln(n_{i,t}) \sim \text{Normal}(\ln(N_i(t)), \sigma_m)$$

where $\sigma_m$ accounts for measurement variation.

For binary values (detected or undetected), we integrated over the possible ranges of viral load values using the same log-normal distribution for the numeric values. For an undetected measurement $(-)$, we integrated over the viral load range $(-\infty, \ln(L)]$, where $L$ is the lower limit of detection of the PCR assay. Similarly, for a detected measurement $(+)$, we integrated over the viral load range $[\ln(L), \infty)$. Lawler et al. (2006) was the only study included in our analysis that reported a lower limit of detection for their PCR assay. For each of the remaining studies, we used the smallest reported viral load value from the study as the effective limit of detection.

To summarize, the likelihood for a set of parameters $\boldsymbol{\theta}$ given data point $n_{i,t}$ and given that individual $i$ was infected is

$$\mathcal{L}_{\text{inf}}(\boldsymbol{\theta}; n_{i,t}) = \begin{cases} f_i(\ln(n_{i,t})) & \text{if } n_{i,t} \text{ is numeric} \\ \int_{\ln(L)}^{\infty} f_i(\ln(n))\, d(\ln(n)) & \text{if } n_{i,t} \text{ is ``detected'' } (+) \\ \int_{-\infty}^{\ln(L)} f_i(\ln(n))\, d(\ln(n)) & \text{if } n_{i,t} \text{ is ``not detected'' } (-) \end{cases}$$

where $f_i(\cdot)$ is the normal density for numeric log viral load values for individual $i$.

##### 1.2.2 Likelihood for Unsuccessful Infections

Given that the individual was not successfully infected, the probability of observing a numeric or "detected" $(+)$ value is $p_{\text{false}}$, which is the false positive probability. This parameter captures the unlikely technical errors that might occur during the course of the experiments. If the measured value was "not detected" $(-)$, then the probability of observing the measurement (a true negative) is $1 - p_{\text{false}}$.

The likelihood for a set of parameters $\boldsymbol{\theta}$ given data point $n_{i,t}$ and given that individual $i$ was non-infected is

$$\mathcal{L}_{\text{noninf}}(\boldsymbol{\theta}; n_{i,t}) = \begin{cases} p_{\text{false}} & \text{if } n_{i,t} \text{ is numeric or ``detected'' } (+) \\ 1 - p_{\text{false}} & \text{if } n_{i,t} \text{ is ``not detected'' } (-) \end{cases}$$

##### 1.2.3 Overall Likelihood

Each individual $i$ has a probability of infection given by the exponential dose-response equation, $P(V_{0,i}) = 1 - \exp(-kV_{0,i})$, where $k$ is the hit probability and $V_{0,i}$ is the initial inoculation dose for individual $i$. The overall likelihood for a set of parameters $\boldsymbol{\theta}$ given a dataset $D$ (combining across all individuals $i$ and times $t$) is

$$\mathcal{L}(\boldsymbol{\theta}; D) = \prod_i \left( P(V_{0,i}) \prod_{t \in T_i} \mathcal{L}_{\text{inf}}(\boldsymbol{\theta}; n_{i,t}) + (1 - P(V_{0,i})) \prod_{t \in T_i} \mathcal{L}_{\text{noninf}}(\boldsymbol{\theta}; n_{i,t}) \right)$$

#### 1.3 Selection and Justification of Prior Distributions

In the following sections we describe the prior distributions for all model parameters. Overall, we sought to set "weakly informative" prior distributions for our model parameters. The goal was to rule out biologically implausible parameter values while at the same time allowing a wide range of biologically plausible parameter values. See **S4 Fig** for prior and posterior predictive simulations.

##### 1.3.1 Infectious Dose and Hit Probability

For improved interpretability, we placed a prior on the infectious dose 50 ($\text{ID}_{50}$) rather than the hit probability ($k$). The hit probability can be computed from the $\text{ID}_{50}$ via $k = \ln(2)/\text{ID}_{50}$. We placed a log-normal prior on the $\text{ID}_{50}$, with the $\text{ID}_{50}$ in units of $\text{TCID}_{50}$:

$$\ln(\text{ID}_{50}) \sim \text{Normal}(\ln(10^4), \ln(10^2))$$

This is a broad prior that places approximately 95% of the probability mass in the range $10^0$ to $10^8$ $\text{TCID}_{50}$. For additional numerical stability, we placed a lower bound of $10^{-10}$ on the hit probability, which corresponds to an $ID_{50}$ of $6.9 \times 10^9$. This is an extremely large and biologically implausible value for the $ID_{50}$, so there is no reason to explore regions of the parameter space with hit probability less than $10^{-10}$.

##### 1.3.2 Viral Kinetics Parameters

For improved interpretability, we placed priors on the doubling time ($t_2$) and halving time ($t_{\frac{1}{2}}$) of the viral load kinetics rather than the growth rate ($g$) and decay rate ($d$). The growth and decay rates can be computed via $g = \ln(2)/t_2$ and $d = \ln(2)/t_{\frac{1}{2}}$. The units for $t_2$ and $t_{\frac{1}{2}}$ are days.

To account for the fact that not all studies used the same strains of SARS-CoV-1 and macaque species, we added study-level hierarchy to these parameters, where $t_{2s}$ and $t_{\frac{1}{2}s}$ are the doubling and halving times for study $s$. The log doubling times are distributed around $\mu_{t2}$ with standard deviation $\sigma_{t2}$, and the log halving times are distributed around $\mu_{t\frac{1}{2}}$ with standard deviation $\sigma_{t\frac{1}{2}}$:

$$\begin{aligned}
\ln(t_{2s}) &\sim \text{Normal}(\mu_{t2}, \sigma_{t2}) \\
\mu_{t2} &\sim \text{Normal}(\ln(0.5), 0.5) \\
\sigma_{t2} &\sim \text{PosNormal}(0, 0.35) \\
\ln(t_{\frac{1}{2}s}) &\sim \text{Normal}(\mu_{t\frac{1}{2}}, \sigma_{t\frac{1}{2}}) \\
\mu_{t\frac{1}{2}} &\sim \text{Normal}(\ln(2), 0.75) \\
\sigma_{t\frac{1}{2}} &\sim \text{PosNormal}(0, 0.35)
\end{aligned}$$

where PosNormal(mode, standard deviation) is a positive-constrained normal distribution with a lower limit of zero. We set broad priors for the mean doubling time ($\mu_{t2}$) and mean halving time ($\mu_{t\frac{1}{2}}$). Approximately 95% of the prior probability mass for $\mu_{t2}$ is in the range 0.18 to 1.36 days, and approximately 95% of the prior probability mass for $\mu_{t\frac{1}{2}}$ is in the range 0.45 to 8.96 days. We expected the growth rate to be higher than the decay rate, so we centered the prior for the mean doubling time on a smaller value than the prior for the mean halving time. We allowed the possibility of studies having highly varied doubling and halving times by setting broad priors for $\sigma_{t2}$ and $\sigma_{t\frac{1}{2}}$.

##### 1.3.3 Dose-Dependence Parameters

For the initial viral load slope parameter ($\alpha_N$) we placed a prior on the difference between the log initial viral load for $10^7$ and $10^5$ $TCID_{50}$ inoculated individuals, $\ln(N_{0,10^7}) - \ln(N_{0,10^5})$. Similarly, for the time of peak viral load slope parameter ($\alpha_t$) we placed a prior on the difference between the log peak time for $10^7$ and $10^5$ $TCID_{50}$ inoculated individuals, $\ln(t_{p,10^7}) - \ln(t_{p,10^5})$. The priors were:

$$\begin{aligned}
\ln(N_{0,10^7}) - \ln(N_{0,10^5}) &\sim \text{Normal}(\ln(100), 1.5) \\
\ln(t_{p,10^7}) - \ln(t_{p,10^5}) &\sim \text{Normal}(0, 1)
\end{aligned}$$

We chose to set priors on these differences rather than directly on the slope parameters for improved interpretability. The prior for the initial viral load difference is centered on ln(100), which corresponds to a 100-fold increase in initial viral load when the inoculation dose is increased from $10^5$ to $10^7$ $TCID_{50}$. We set a wide prior for the initial viral load difference, with approximately 95% of the prior probability mass for the ratio of initial viral load between the $10^7$ and $10^5$ $TCID_{50}$ doses in the range $[4.98, 2008.55]$. The prior for the time of peak viral load difference is centered on 0, which corresponds to the scenario where the inoculation dose does not affect the time of peak viral load. We set a wide prior for the time of peak viral load difference, with approximately 95% of the prior probability mass for the ratio of time of peak viral load between the $10^7$ and $10^5$ $TCID_{50}$ doses in the range $[0.14, 7.39]$. The slope parameters can be calculated from these log differences as

$$\begin{aligned}
\alpha_N &= \frac{\ln(N_{0,10^7}) - \ln(N_{0,10^5})}{\ln(\bar{V}_{s,10^7}) - \ln(\bar{V}_{s,10^5})} \\
\alpha_t &= \frac{\ln(t_{p,10^7}) - \ln(t_{p,10^5})}{\ln(\bar{V}_{s,10^7}) - \ln(\bar{V}_{s,10^5})}
\end{aligned}$$

where $\bar{V}_{s,10^7}$ and $\bar{V}_{s,10^5}$ correspond to the expected number of successful virions given an inoculation dose of $10^7$ and $10^5$ $TCID_{50}$, respectively.

We placed normal priors on the intercept parameters, $\beta_N$ and $\beta_t$. To account for between-study variation (e.g., one study tends to measure higher viral loads than other studies), we added study-level hierarchy to these parameters, where $\beta_{Ns}$ and $\beta_{ts}$ are the intercept parameters for study $s$. The log initial viral load intercept parameters are distributed around $\mu_{\beta N}$ with standard deviation $\sigma_{\beta N}$, and the log time of peak viral load intercept parameters are distributed around $\mu_{\beta t}$ with standard deviation $\sigma_{\beta t}$:

$$\begin{aligned}\beta_{Ns} &\sim \text{Normal}(\mu_{\beta N}, \sigma_{\beta N})\\ \mu_{\beta N} &\sim \text{Normal}(\ln(10^3), \ln(10))\\ \sigma_{\beta N} &\sim \text{PosNormal}(0, 0.35)\\ \beta_{ts} &\sim \text{Normal}(\mu_{\beta t}, \sigma_{\beta t})\\ \mu_{\beta t} &\sim \text{Normal}(\ln(2), 0.5)\\ \sigma_{\beta t} &\sim \text{PosNormal}(0, 0.35)\end{aligned}$$

We set broad priors for the intercept parameters. The parameter $\mu_{\beta N}$ is the mean for the initial viral load intercept parameters, which are the log initial viral loads given a successful infection following an inoculation dose of $10^5$ $TCID_{50}$. Approximately 95% of the prior probability mass for the mean of the initial viral load intercept parameter falls in the range 10 to $10^5$ RNA copies per sample. The parameter $\mu_{\beta t}$ is the mean for the time of peak viral load intercept parameters, which are the log times of peak viral load given a successful infection following an inoculation dose of $10^5$ $TCID_{50}$. Approximately 95% of the prior probability mass for the mean of the time of peak viral load intercept parameter falls in the range 0.74 to 5.44 dpi. We allowed the possibility of studies having highly varied intercept parameters by setting broad priors for $\sigma_{\beta N}$ and $\sigma_{\beta t}$.

##### 1.3.4 Measurement Variation

We placed log-normal priors on the measurement variation parameters, $\sigma_m$. To account for between-study variation, we added study-level hierarchy to these parameters, where $\sigma_{ms}$ is the measurement variation for study $s$:

$$\ln(\sigma_{ms}) \sim \text{Normal}(\ln(1), \ln(1.5))$$

This is a broad prior that places approximately 95% of the prior probability mass for the measurement variation ($\sigma_m$) in the range $[0.44, 2.25]$.

##### 1.3.5 False Positive Probability

We placed a log-normal prior on the false positive probability:

$$\ln(p_{\text{false}}) \sim \text{NegNormal}(\ln(10^{-4}), \ln(1.5))$$

where NegNormal(mode, standard deviation) is a negative-constrained normal distribution with an upper limit of zero. The prior for the false positive probability is centered on a small value ($10^{-4}$) with a relatively narrow distribution, which encodes our assumption that false positives are rare in these experiments. We used a negative normal distribution to ensure that $\ln(p_{\text{false}}) \leq 0$ and $p_{\text{false}} \in [0, 1]$.
