## Supplementary figures and images for "Data aggregation and mechanistic modeling enable dose-response analysis of SARS-CoV-1 in non-human primates"

### Fig S1

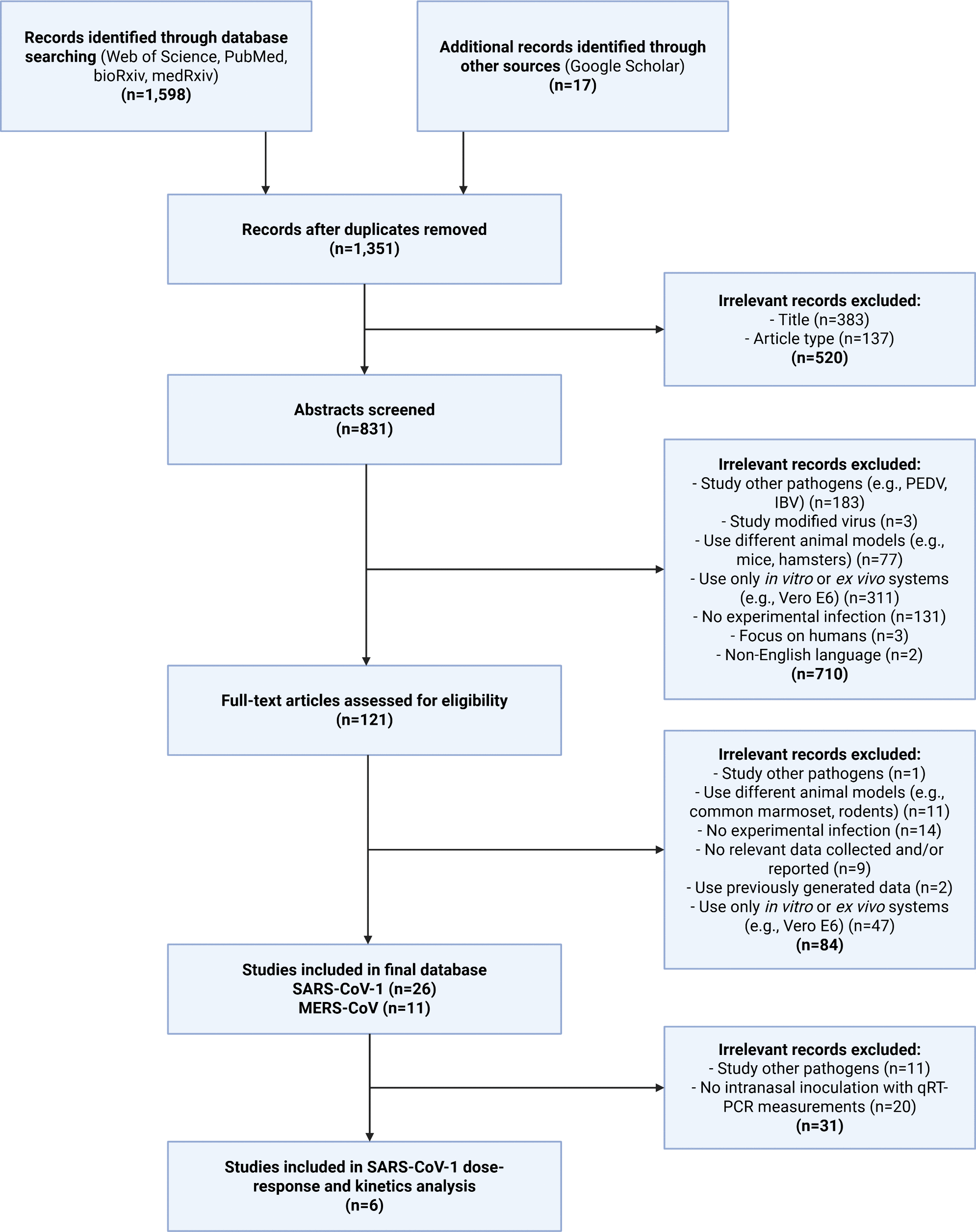

### Fig S2

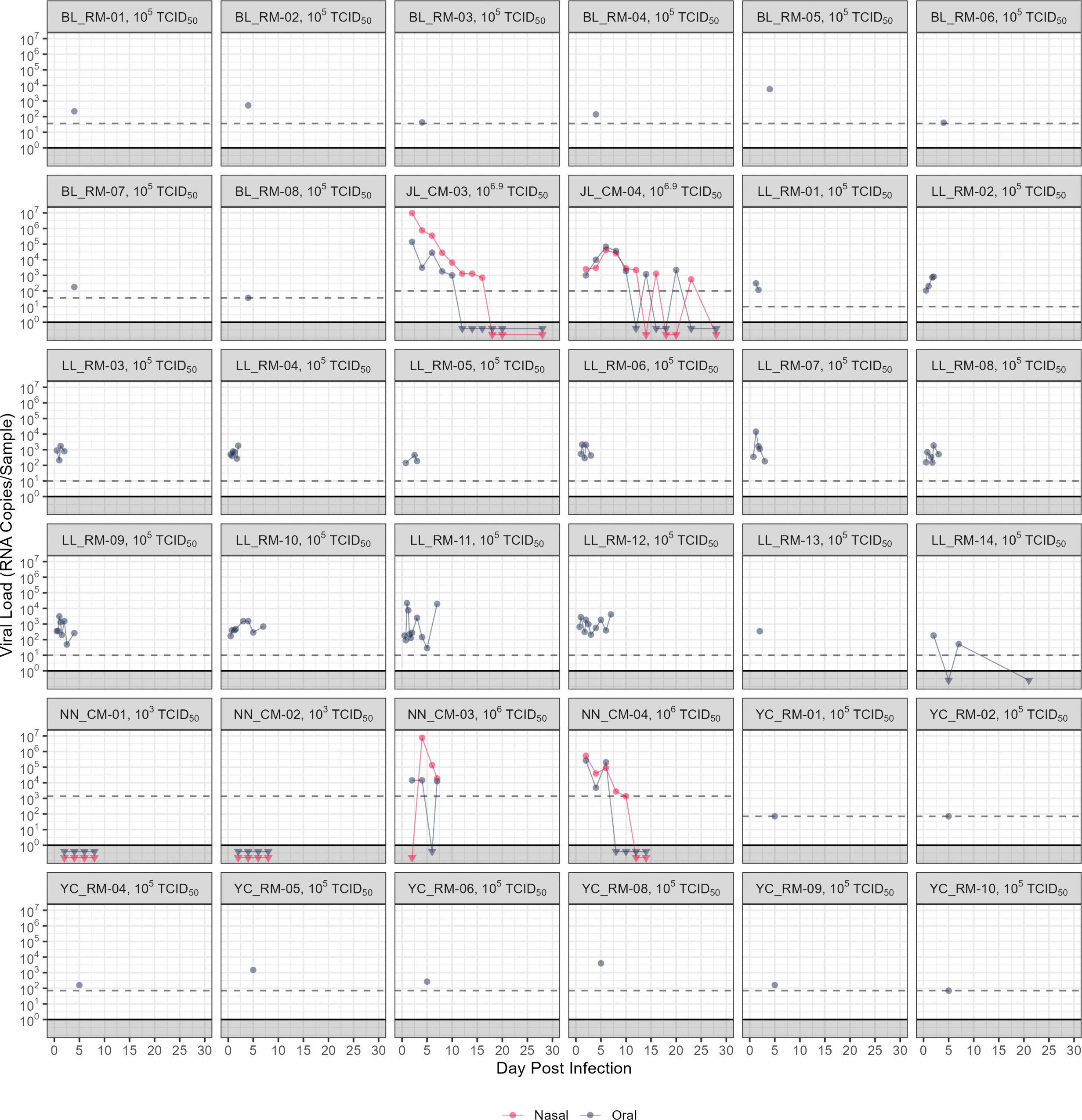

### Fig S3

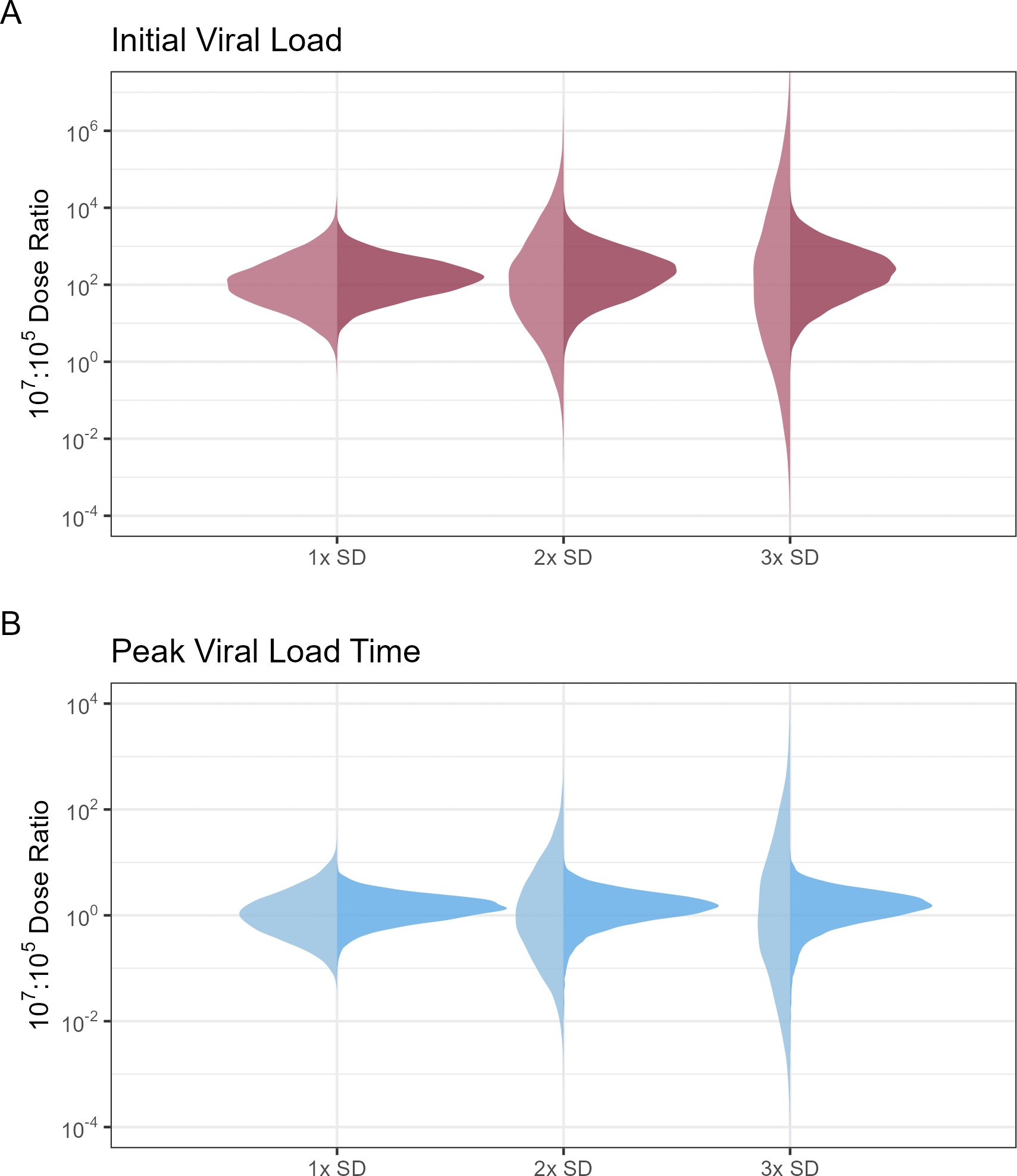

### Fig S4

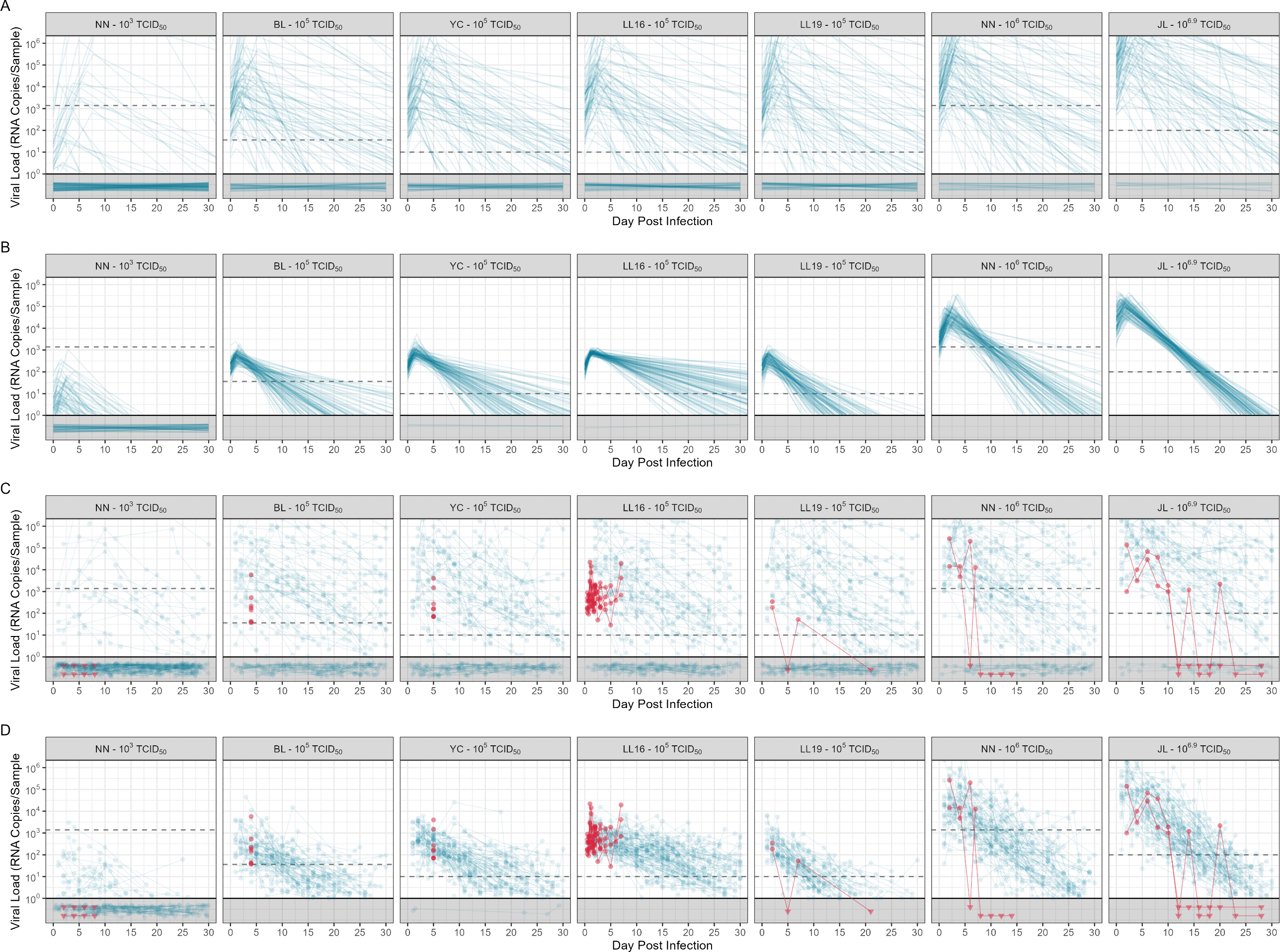

### Fig S5

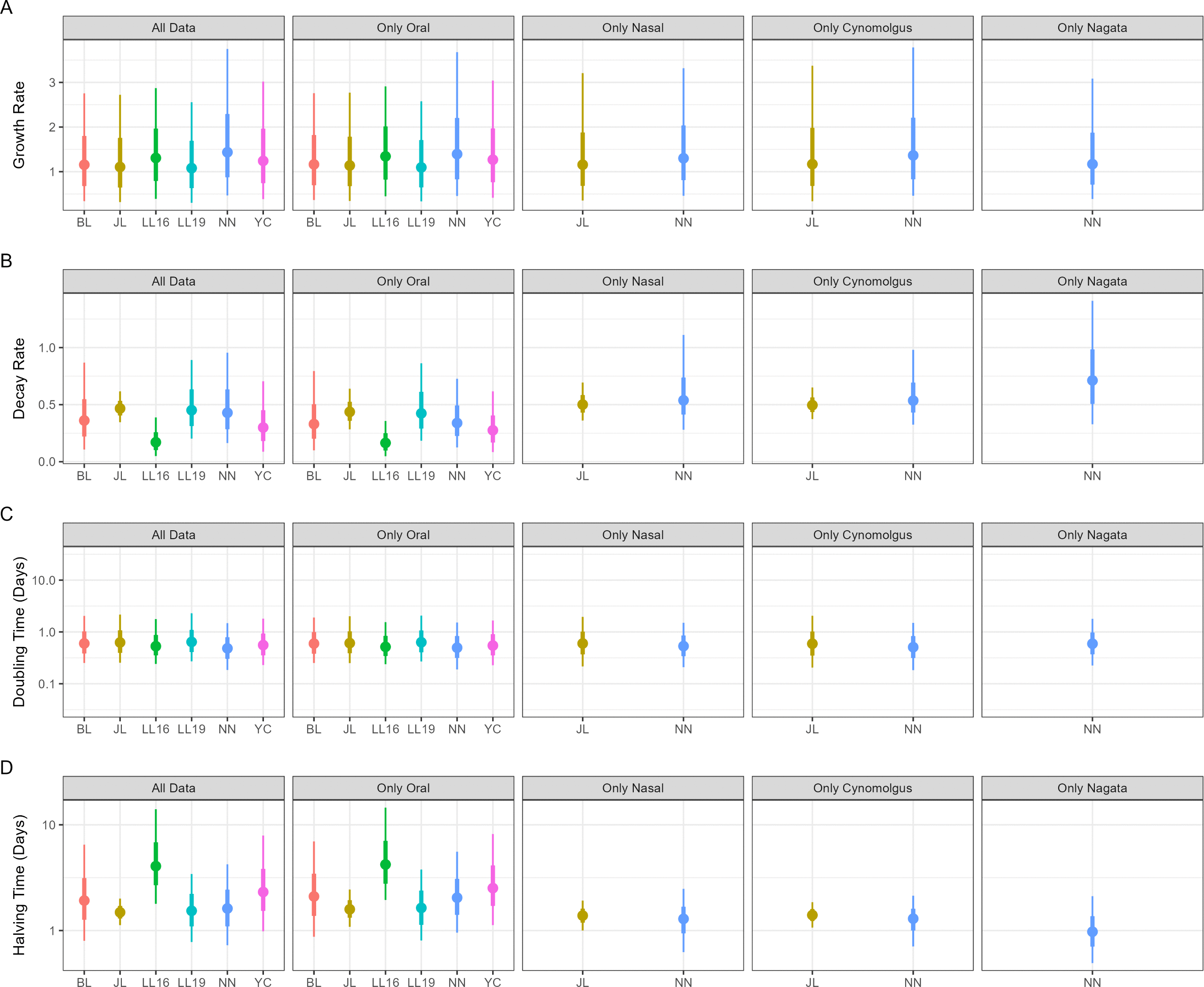

### Fig S6

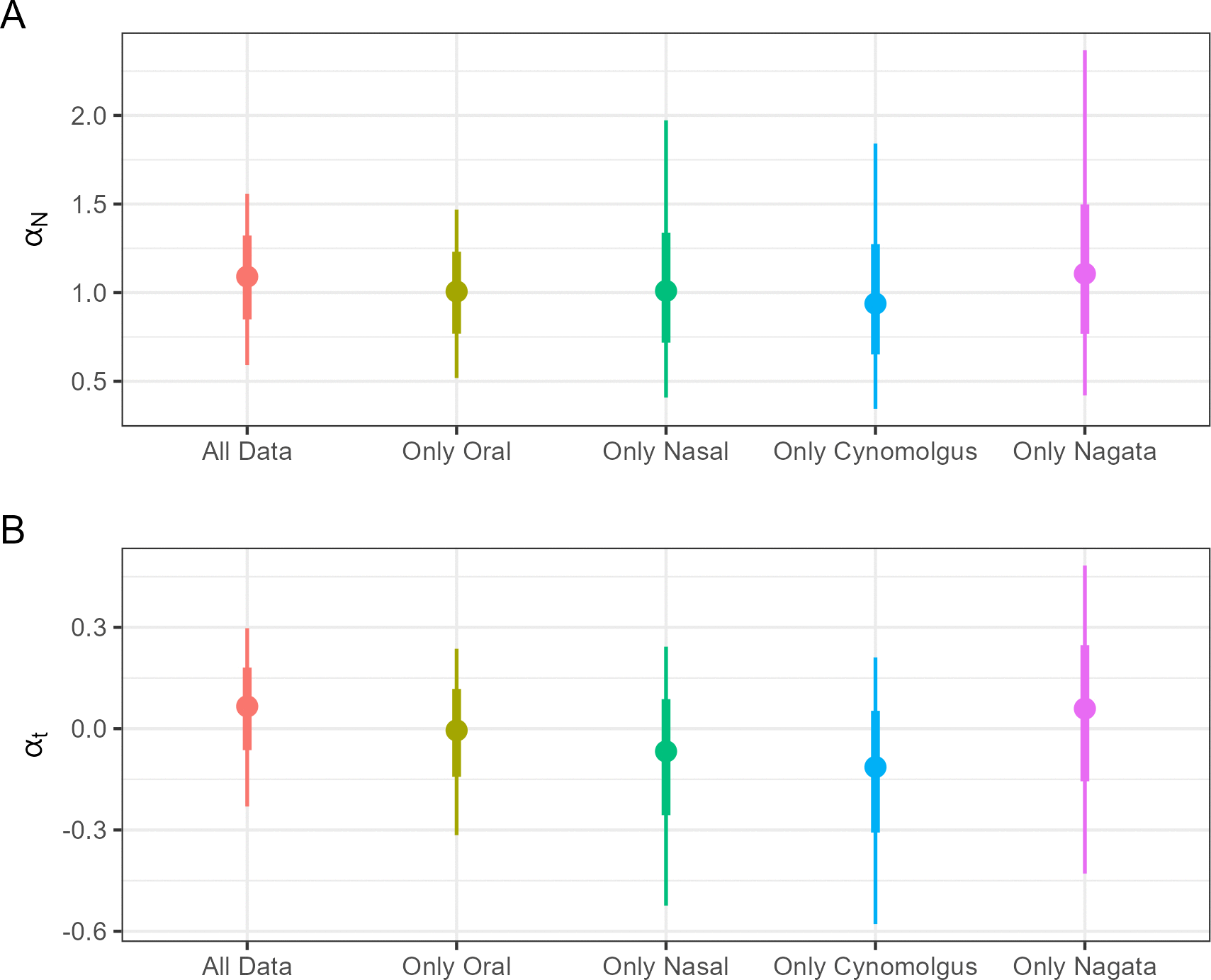

### Fig S7

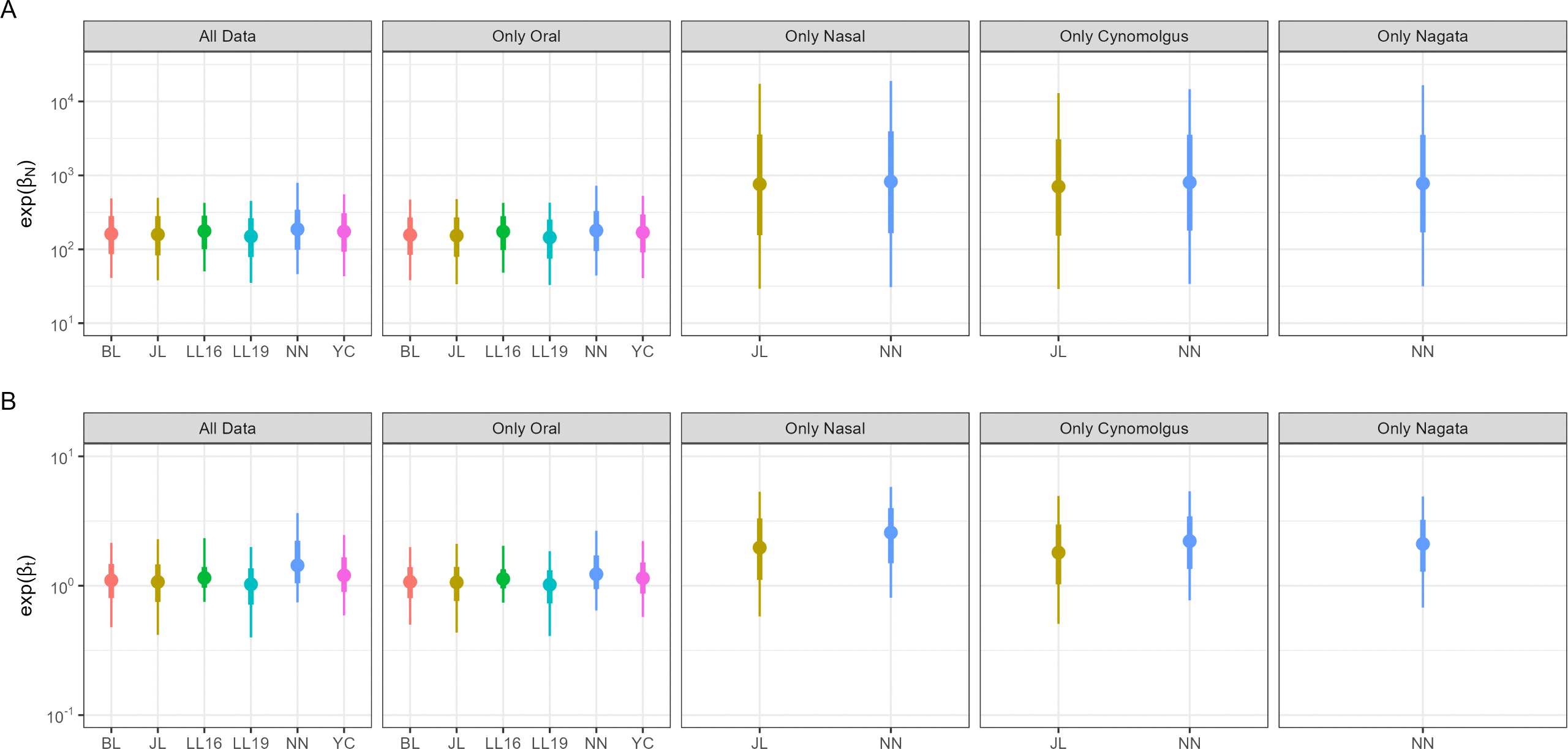
